# MRI-Based Structural Development of the Human Newborn Hypothalamus

**DOI:** 10.1101/2025.06.20.660741

**Authors:** Elizabeth Yen, Josepheen DeAsis-Cruz, Jerod M. Rasmussen

## Abstract

**Background:** Preclinical evidence suggests that intrauterine exposures can impact hypothalamic structure at birth and future disease risk, yet early human data are limited.

**Methods:** Hypothalamic volumes were measured from 699 T1-weighted MRI scans from 631 newborns (54% female; 27–45 weeks postmenstrual age/PMA) in the Developing Human Connectome Project. Linear mixed-effects models tested associations with prenatal exposures: gestational age (GA) at birth, PMA at scan, sex, maternal body mass index (BMI), and smoking. Findings were partially replicated in the Adolescent Brain and Child Development (ABCD) Study (release 5.1) data (16,934 observations from 11,207 participants).

**Results:** Absolute hypothalamus volume increased with PMA (+5.5%/week, t=39.9, p<10⁻¹⁰), but not after adjusting for brain volume (t=1.2, p=0.24). Males showed larger absolute (+3.3%, t=3.2, p=0.002) but smaller relative hypothalamus volume (t=-2.8, p=0.005). Lower GA was linked to larger relative hypothalamus volume (t=-6.5, p<10⁻⁹), with evidence for sex moderation (t=-2.4, p=0.019). Smoking during pregnancy was associated with smaller hypothalamus volume in newborns (t=-2.05, p=0.04; dose dependence: t=-2.9, p<0.01). Smoking remained associated with reduced hypothalamus volume in adolescents (t=-2.8, p=0.005).

**Conclusions:** The findings suggest that the hypothalamus is a crucial and underexplored target of perinatal influences for understanding the origins of long-term health and disease.

**Impact:**

- This study highlights the hypothalamus as a critical and underexplored target for understanding how prenatal exposures in human newborns could influence long-term health and disease.
- Gestational age (GA) at birth, postmenstrual age at scan, and smoking during pregnancy are associated with hypothalamic volume in newborns.
- The effects of GA on adjusted hypothalamic volume appear to be transient, while the effects of smoking seem to last throughout adolescence.
- Our findings suggested sex-specific effects on the associations between volume and age measures across development.

## 1. Background

Recent advances in brain magnetic resonance imaging (MRI) continue to enhance our understanding of the developing fetal and newborn brain (1–5). Specifically, this line of research has begun to uncover how maternal health conditions during pregnancy affect the developing brain, with long-term consequences for a range of outcomes, including physical growth, cognitive function, school performance, and psychiatric illness (6–9). Much of this attention has focused on global brain development and the cortical and subcortical systems that support higher-order functions (10). While this attention is warranted, considerably less focus has been directed to a “small but mighty” (11) brain structure essential for sustaining life through the regulation of physiological homeostasis and control of survival-oriented behaviors: the hypothalamus.

The hypothalamus is a highly conserved structure across vertebrate species (12,13) and controls fundamental functions, including thermoregulation, wake-sleep cycles, energy metabolism, fluid and electrolyte balance, feeding and suckling, drinking, and reproduction (14). It is among the earliest regions of the central nervous system to develop (15), emerging as early as the third week of gestation in the form of the neural plate (16), making it especially vulnerable to in-utero exposures that shape development in anticipation of postnatal conditions. The hypothalamus plays a unique and critical role in fetal and early postnatal life due to its exponential growth and development during this period (17). Cross-species studies demonstrate the importance of optimal hypothalamus development during gestation for a wide range of behavioral and neuropsychological adaptations to ensure the species’ ability to deal with early life stressors, which in turn determine the physiological and psychological health outcomes later in life (18,19). For example, maternal and pregnancy-related inflammatory conditions may disrupt normal hypothalamic development, leading to early behavioral changes that predispose offspring to adult-onset diseases (20–22). However, its small size (<1 cm^³^ in adults) (23) and the lack of tools available for its delineation have constrained progress in understanding the developing hypothalamus, with recent notable exceptions (24,25). Thus, given its critical role in physiological regulation, the hypothalamus remains an essential but underexplored target for advancing understanding of the developmental origins of health and disease.

A recently developed registration-based pipeline enables FreeSurfer-compliant automated segmentation of the neonatal hypothalamus and its subunits, validated on over 200 brain MRIs across four NIH Environmental Influences on Child Health Outcomes (ECHO) sites (26). In the present study, this infant-optimized pipeline was applied to a large publicly available newborn dataset (Developing Human Connectome Project, dHCP, United Kingdom) to investigate perinatal influences (gestational age at birth [GA], maternal pre-pregnancy body mass index [BMI], maternal smoking during pregnancy) on newborn hypothalamic development. Because the time near birth represents a period largely antecedent to postnatal exposures, this design maximizes outcome variance attributable to prenatal exposures (27). To test whether maternal factors during pregnancy have lasting effects on hypothalamic development following postnatal exposures, we leverage a second large, publicly available adolescent dataset (Adolescent Brain and Child Development, ABCD, United States)(28). Finally, given the widely recognized sexual dimorphism in brain development (29,30), we also evaluate sex differences in response to perinatal influences.

## 2. Methods

### 2.1 Data collection

Neuroimaging (MRI) and associated data were obtained from the Developing Human Connectome Project (dHCP) study, located in the NIMH Data Archive (Collection #3955). This dataset includes a large number of healthy, term-born, and preterm infants, allowing for the characterization of typical development. Additional details about this cohort are well described elsewhere (31,32). For the current study, 699 observations of 631 unique participants were analyzed. Infants born preterm were included to increase variation in postmenstrual age (PMA) at scan, thereby improving the precision of estimated age-related slopes. Demographic and exposure characteristics are described in **Table 1**. Pregnant women were enrolled from mid to late pregnancy, and imaging occurred between 26.7 and 45.1 weeks PMA (**Figure 1**). Exclusion criteria were contraindication to MR imaging, preterm infants deemed clinically unstable to tolerate scanning (e.g., thermoregulation issues, sepsis, need for invasive mechanical ventilation), and language barriers impacting the process. Data collection and dissemination were approved by the United Kingdom Health Research Authority (Research Ethics Committee reference number: 14/LO/1169). In addition, a partial replication analysis was used to test the persistence of effects into adolescence using the ABCD Study as approved by a central Institutional Review Board. For a more complete description of recruitment and inclusion criteria, we refer the reader to articles by (28,33). Written parental consent was obtained in all cases.

**Figure 1.**
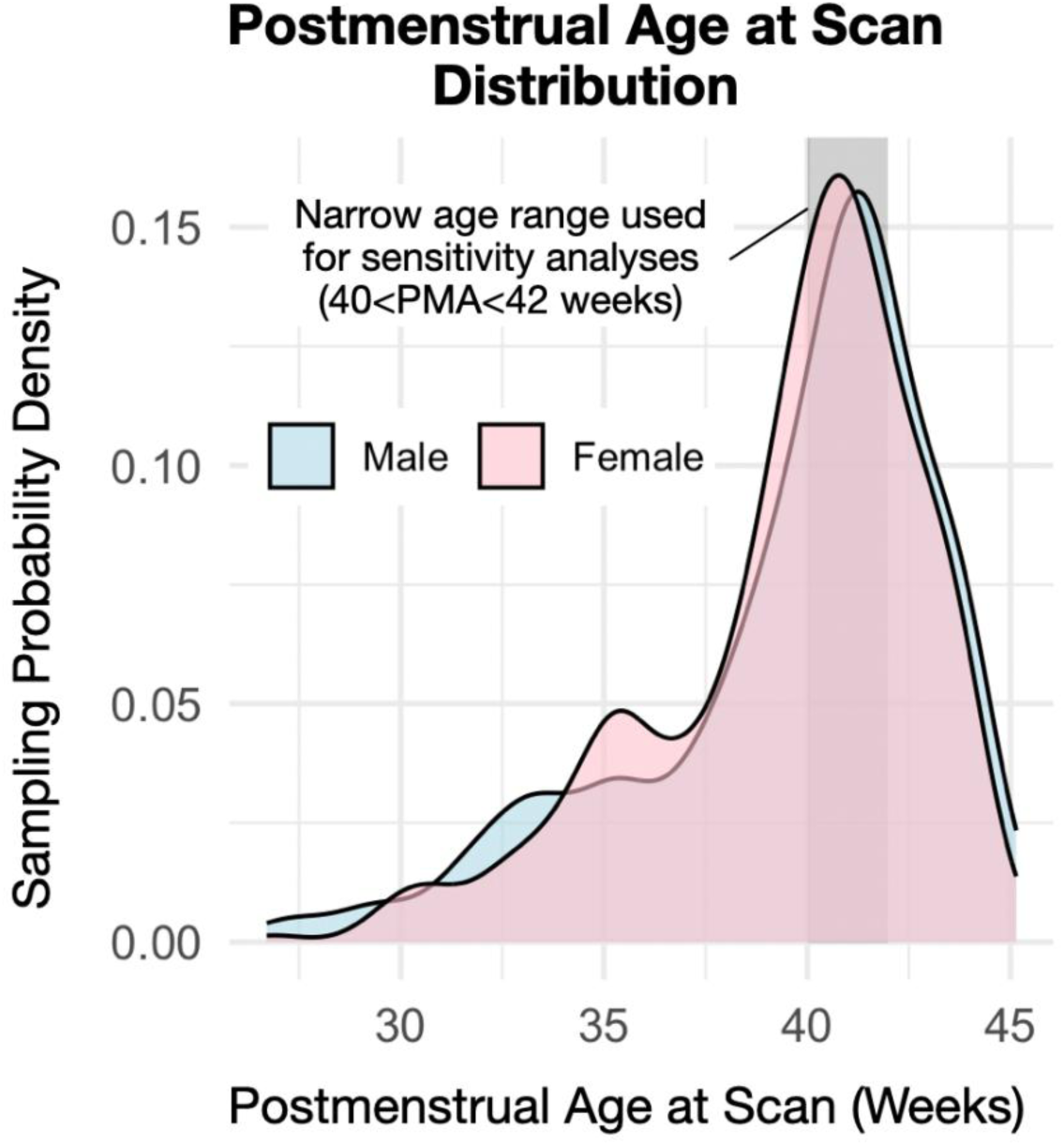
Postmenstrual age at scan distribution by sex. No significant sex differences were observed in PMA at scan, GA, maternal smoking status, or maternal pre-pregnancy BMI.

**Table 1.**
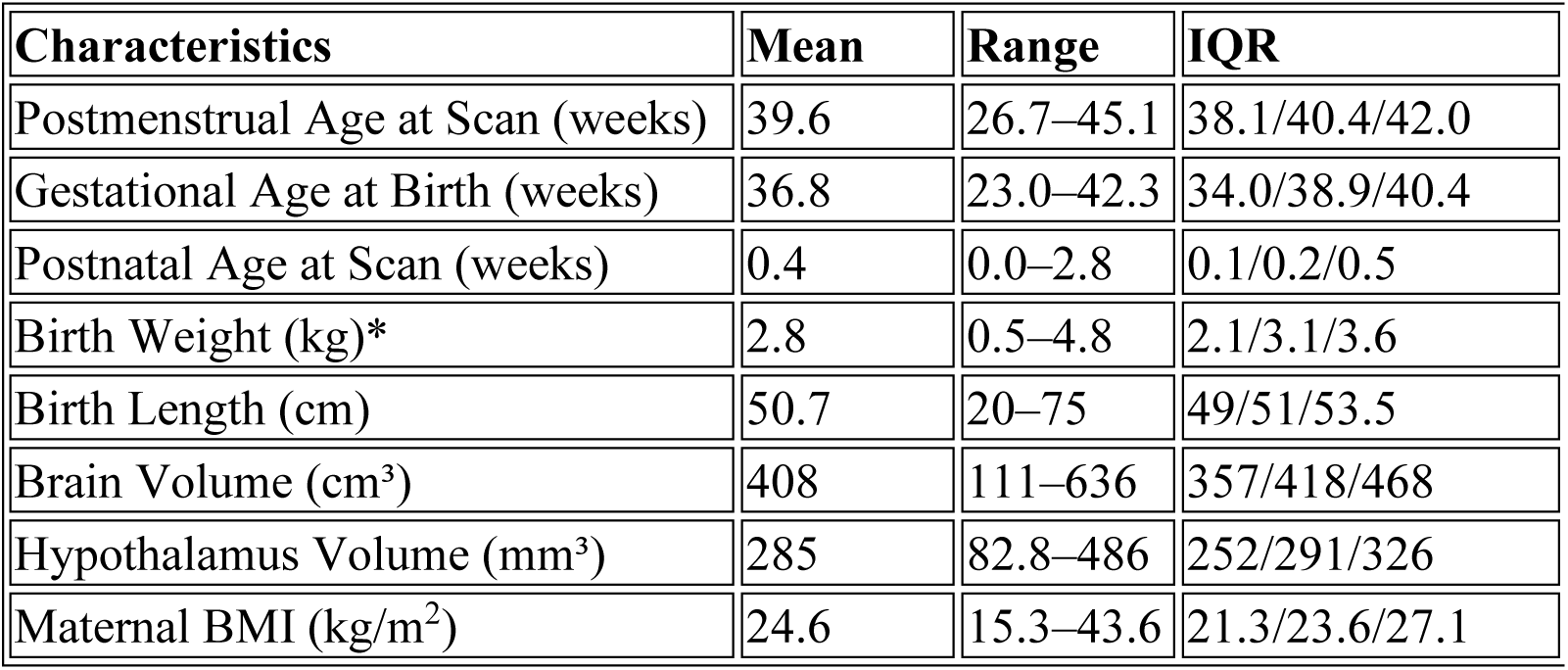
Study Demographics. The newborn participants were split 53.6% female/46.4% male. Maternal smoking during pregnancy was recorded in 3.3% of participating mothers. *Males were ∼75g larger in birth weight than their female counterparts after adjustment for GA (p=10^-4^). No other sex differences in table demographics were observed (all p>0.4) except for brain and hypothalamus volume (discussed in detail below).

| Characteristics | Mean | Range | IQR |
| --- | --- | --- | --- |
| Postmenstrual Age at Scan (weeks) | 39.6 | 26.7–45.1 | 38.1/40.4/42.0 |
| Gestational Age at Birth (weeks) | 36.8 | 23.0–42.3 | 34.0/38.9/40.4 |
| Postnatal Age at Scan (weeks) | 0.4 | 0.0–2.8 | 0.1/0.2/0.5 |
| Birth Weight (kg)* | 2.8 | 0.5–4.8 | 2.1/3.1/3.6 |
| Birth Length (cm) | 50.7 | 20–75 | 49/51/53.5 |
| Brain Volume (cm<sup>3</sup>) | 408 | 111–636 | 357/418/468 |
| Hypothalamus Volume (mm<sup>3</sup>) | 285 | 82.8–486 | 252/291/326 |
| Maternal BMI (kg/m<sup>2</sup>) | 24.6 | 15.3–43.6 | 21.3/23.6/27.1 |

T1-weighted images were used to segment the hypothalamus (see *Infant Hypothalamus Segmentation* below). For the dHCP study, MRI was performed at the Evelina London Children’s Hospital on a 3T Philips Achieva system with a dedicated 32-channel neonatal head coil. T1- weighted inversion recovery images were collected as part of a newborn imaging battery (TR=4795 ms; TI=1740 ms; TE=8.7 ms; SENSE factor: axial=2.26, sagittal=2.66). One image stack each (5m45s) in sagittal and axial slice orientation was acquired with 0.8mm in-plane resolution and 1.6 mm slices overlapped by 0.8 mm. Images were reconstructed at a 0.8mm isotropic resolution and downloaded for use here as provided by the dHCP data repository.

### 2.2 Infant Hypothalamus Segmentation

Infant hypothalamic volumes were derived using an adapted version of the recently published and publicly available (https://github.com/jerodras/neonate_hypothalamus_seg) segmentation pipeline, segATLAS (26). In brief, this method uses a FreeSurfer-compatible hypothalamus atlas in high-quality PMA-appropriate template space (Baby Connectome Project, BCP) (34) and robust infant brain extraction methods (ANTsPyNet, https://github.com/ANTsX/ANTsPyNet) consistent with the BCP template definitions (12). However, given the high-quality image registrations publicly shared by dHCP and their now ubiquitous community use, we integrated the existing dHCP registrations to the 40-week Serag template into the existing segATLAS pipeline. This was done by creating an atlas in the 40-week Serag template space using ANTs non-linear registration. Estimates of volume were then based on a soft-segmentation approach, adding up fully- and partially-volumed voxels. Similarly, brain volume (BV) is based on the dHCP-provided brain masks and used here to reflect global brain size, controlling for non-specific hypothalamic growth. In addition, to test the specificity of effects (*i.e.,* not generalizable to other subcortical structures), bilaterally averaged amygdala volumes were chosen as a negative control (21) based on their comparable size to the hypothalamus and relative spatial proximity as a central brain structure.

### 2.3 Modeling Effects of Growth and Prenatal Predictors

As a first step towards understanding hypothalamic growth in early life, we constructed a base model using linear mixed-effects regression to regress hypothalamus volume onto PMA at scan and sex. This base model did not include other prenatal or demographic predictors. Further, to understand hypothalamic growth beyond global increases in brain size, we used two complementary approaches. First, BV was added as a term to the base linear model, allowing for independent effects (i.e., growth of the hypothalamus) beyond global size. Alternatively, hypothalamus volume was expressed as a ratio relative to BV, thereby forcing a relationship between hypothalamus volume and BV and accounting for non-linearity (a ratio approach). The rationale for these two approaches is that while the ratio approach accounts for non-linearities in growth that are not accounted for in linear regression, it does so by forcing BV variance into the outcome. Because growth in this age range is expected to be approximately linear, the non-ratio approach was our first choice while confirming that any observed relationships also hold true using the ratio approach as described in the *Supplementary Material*.

GA, maternal pre-pregnancy BMI, and maternal cigarette smoking during pregnancy were considered as prenatal predictors of early life hypothalamus volume over and above adjustment for PMA, BV, and sex (base model). GA was estimated from the last menstrual period and/or confirmed by early ultrasound. PMA was defined as GA plus chronological age (i.e., the time since birth). Maternal pre-pregnancy BMI and smoking status were collected by survey at the time of enrollment. Maternal smoking status during pregnancy was coded as a dichotomous yes/no answer based on self-report. Additionally, for a dose-response analysis, we used the numeric response to “Average number of cigarettes smoked a day at that time.” A full linear mixed-effects model was constructed by expanding the base model above to include prenatal predictors. Final model selection was performed using a best subset analysis based on adjusted R-squared selection criteria implemented using the “leap” package in R version 4.3.3. Lastly, sex-specific effects were considered by testing for sex interactions whenever feasible.

### 2.4 Partial Replication Analysis in an Independent Adolescent Cohort

To determine if neonatal and maternal factors have a persistent effect on hypothalamic volumes extending into adolescence, we conducted a partial replication analysis assessing the effects of chronological age, sex, GA, and maternal smoking during pregnancy on hypothalamus volume in an independent cohort (ABCD study). High-quality T1-weighted images from 11,207 individuals (16,934 observations, 9,380 unique families) in the ABCD study were considered for analysis (ABCD Data Release 5.1). Imaging data were available at baseline (∼10 years of age) and two-year follow-up. Hypothalamus volume was derived using FreeSurfer’s *mri_segment_hypothalamic_subunits* (35) tool and harmonized by accounting for site effects using ComBat batch correction (36). Age at interview (interview_age), sex (demo_sex_v2), and brain volume (mrisdp_604) were based on ABCD-provided tabulated measures. Information on the exact GA was not available, therefore, preterm birth status (devhx_12a_p, yes/no < 37-weeks GA) was used as a proxy. Maternal smoking status during pregnancy was available for analysis (devhx_9_tobacco, “Once you knew you were pregnant, were you using any of the following: Tobacco?”). As above, a parsimonious linear mixed-effects model considering chronological (adolescent) age, sex, and BV was used to test for an association with hypothalamus volume and nesting for participant and family structure using random intercepts. Informed by the dHCP findings, this model was repeated testing for an age-by-sex interaction effect. A second linear mixed-effects model, adding maternal smoking status during pregnancy and preterm birth status, was used to test for an association with hypothalamus volume over and above age, sex, and brain volume. Lastly, a full mixed-effects model, adding potentially confounding factors (area deprivation index: adi_weighted; parent education: prnt_educ_max; household income: prnt_comb_income_cont; puberty status: ppdms_score; race: race_cleaned; and ethnicity: demo_ethn_v2; 14,446 complete observations across 7,732 individuals), was used to test for an independent association with hypothalamus volume.

## 3. Results

### 3.1 Postmenstrual Age at Scan is Associated with Absolute, but not Relative Hypothalamus Volume

Raw, unadjusted (absolute) hypothalamus volume was strongly associated with PMA (**Figure 2**, +5.5% increase in volume per week, t=39.9, p<10^-10^, 699 observations across 631 unique participants). The normalized slope of the association with PMA is significantly different when compared to that of BV (t=-4.8, p<10^-5^), suggesting that the hypothalamus grows at a slower rate. Evaluation of the amygdala also showed a significantly slower growth rate when compared to BV (t=-3.2, p<0.001). There were no significant differences when comparing the relative hypothalamus to amygdala growth (t=1.5, p=0.12). This suggests that the observed difference in hypothalamus slope is not specific to the hypothalamus, but rather generalizable to at least one other subcortical structure. In addition, hypothalamus volume was not significantly associated with PMA when adjusting for BV in a linear model (t=1.2, p=0.24) or as a ratio (t=-1.0, p=0.32). Hence, while the slope of association with PMA differs between *absolute* hypothalamus volume and BV, these differences were not observed when expressed as *relative* measures.

**Figure 2.**
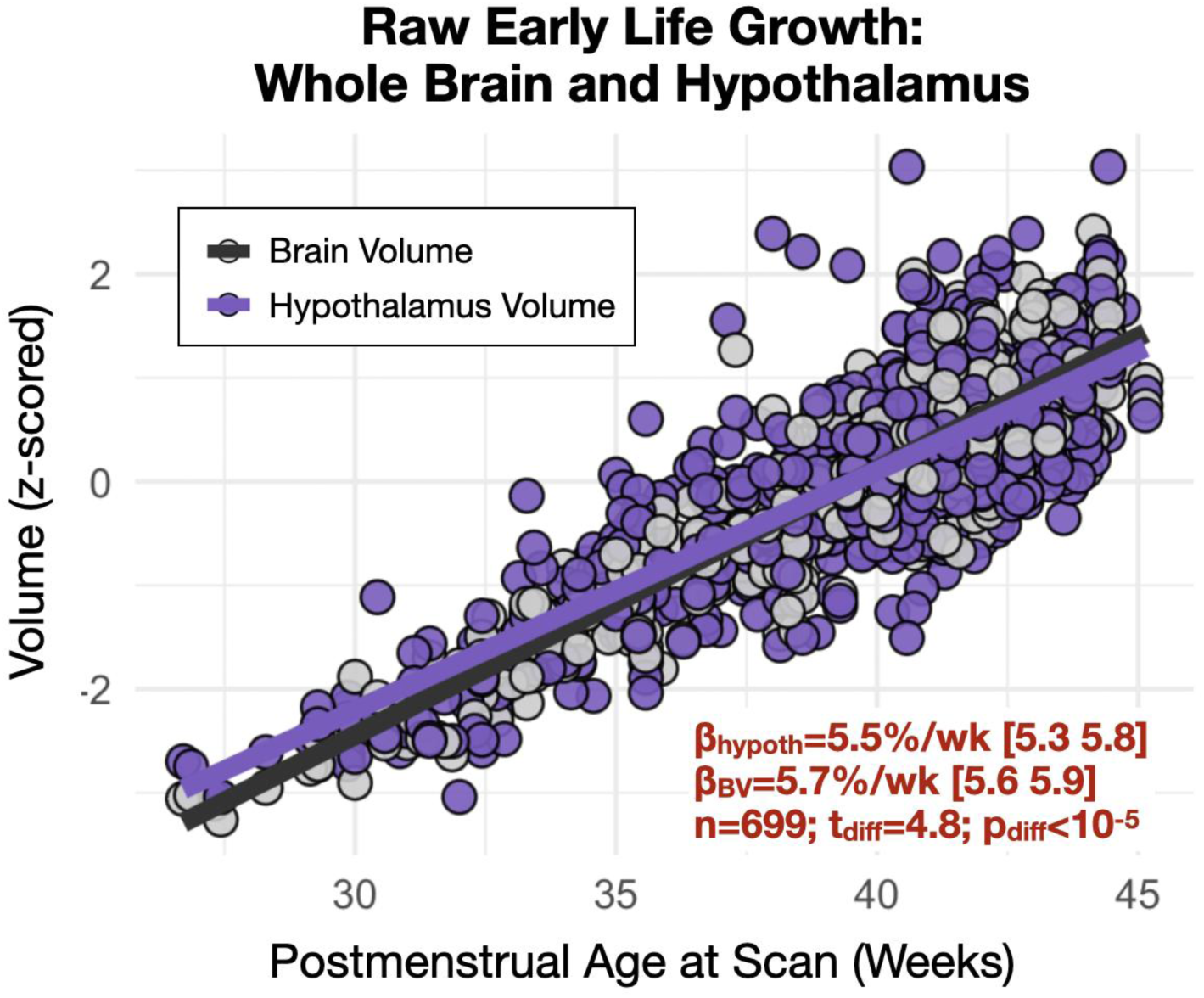
Postmenstrual age at scan is associated with raw (unadjusted) hypothalamus volume. Hypothalamus volume was associated with PMA. While the slope of association with age differs between *absolute* hypothalamus volume and BV (p<10^-5^), no association was observed when expressing hypothalamus volume as a measure *relative* to BV.

### 3.2 Infant Sex is Associated with Absolute and Relative Hypothalamus Volume

In addition to the significant effect of PMA, infant sex also significantly affects BV and hypothalamus volume. Compared to females, males had larger *unadjusted* global brain and hypothalamus volumes (BV: male=+7.0%, t=8.9, p<10^-10^, hypothalamus: male=+3.3%, t=3.2, p=0.002). A significant effect of sex on hypothalamus volume remained in both the adjusted model and the ratio model, with smaller *relative* hypothalamic measures in males compared to females (t=-2.8, p_adj_=0.005; t=-3.7, p_ratio_<10^-5^). This effect was not observed using amygdala volumes (t=0.3, p_ratio_=0.76), suggesting that the observed effects of sex were specific to the hypothalamus.

### 3.3 Gestational Age at Birth and Maternal Smoking During Pregnancy Are Associated with Infant Hypothalamus Volume

GA and maternal smoking during pregnancy were associated with infant hypothalamus volume after adjusting for PMA, BV, and sex (**Figures 3 and 4**, **Table 2**). No association was observed between maternal pre-pregnancy BMI and infant hypothalamus volume (t=-0.5, p=0.69; excluded based on best subsets model fit). The direction of association with GA was negative, suggesting that early birth is consistent with an enlarged hypothalamus relative to BV (**Figure 3**). We also observed sex-specific effects with point estimates suggesting males have larger adjusted relative hypothalamus volumes early in life (∼25 weeks GA) but with a larger negative GA slope compared to females. Maternal smoking (dichotomous yes/no) during pregnancy was associated with a smaller hypothalamus. Notably, we also observed a dose-dependent effect in the subset of mothers (n=23) who reported smoking during pregnancy (t=-2.9, p<0.01) (**Figure 4**). This effect was not observed when using amygdala volume as a comparison (t_yes/no,amyg_=0.23, p_yes/no,amyg_=0.82; t_dose,amyg_=-0.88, p_dose,amyg_=0.32). Model diagnostics for multi-collinearity suggested that while the effects of PMA and BV are potentially inflated in the model, effects of sex, GA and smoking are not (VIF_PMA_=6.6, VIF_BV_=5.9, VIF_GA_=1.7, VIF_sex_=1.1, VIF_smoking_=1.0).

**Figure 3.**
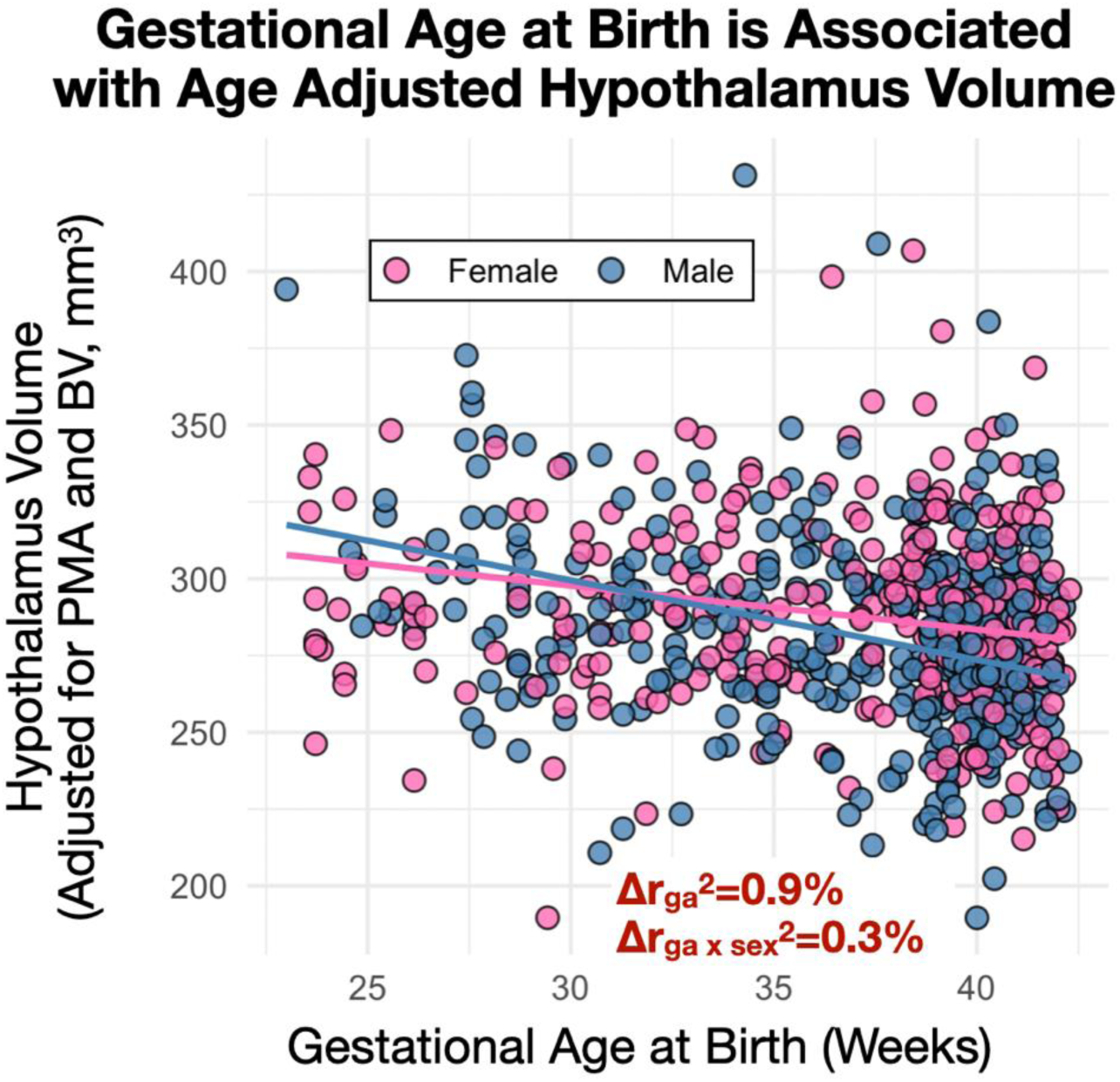
Gestational age and early life hypothalamus volume. a) GA was negatively associated with hypothalamus measures adjusted for PMA and BV (t=-6.5; p<10^-9^) and exhibited sex-specific associations (t=-2.4; β_m,brain_>β_f,brain_: p=0.019).

**Figure 4.**
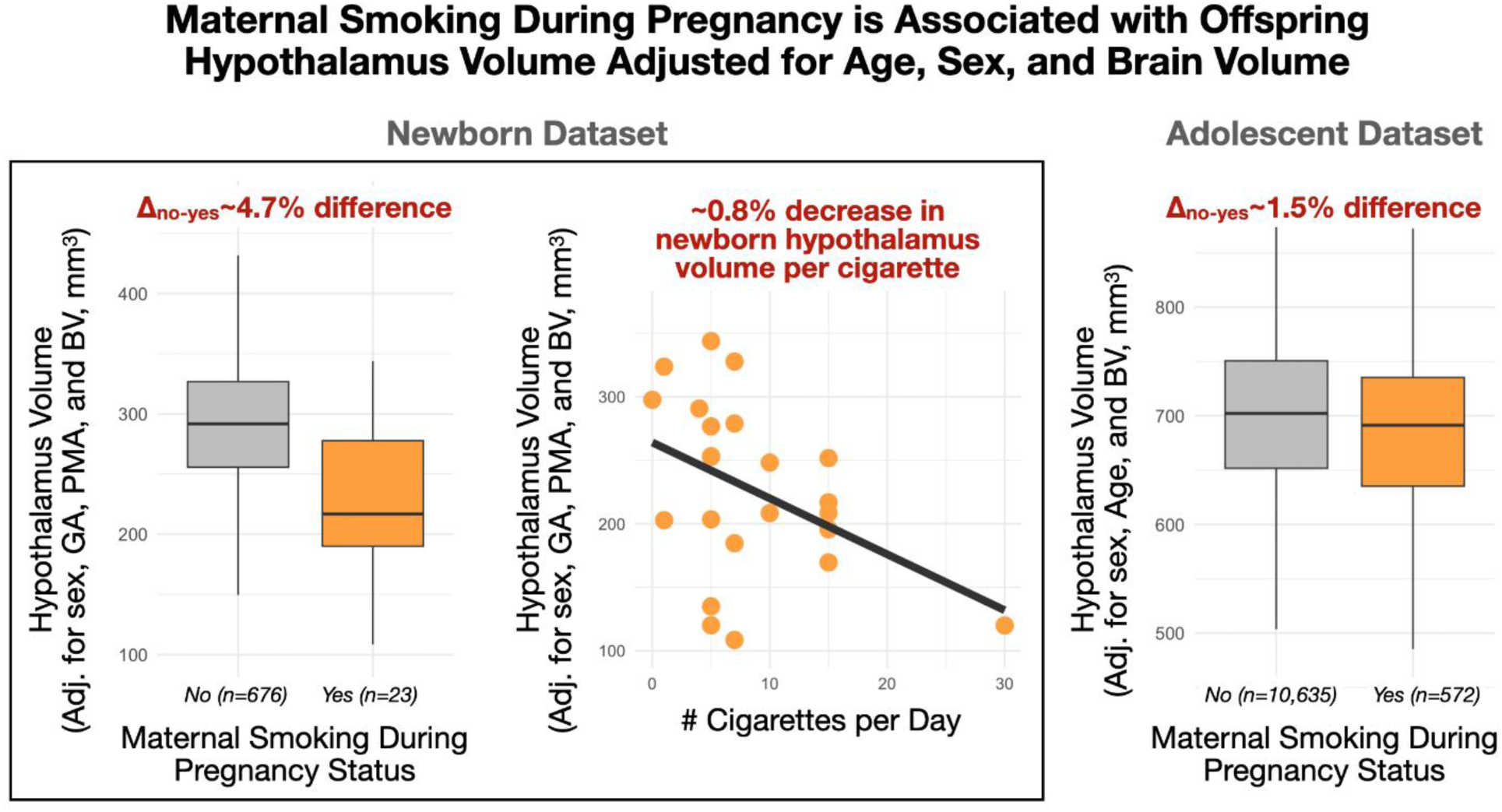
Effect of maternal cigarette smoking status on offspring hypothalamus volume. Offspring born to mothers who smoked during pregnancy exhibited smaller postnatal hypothalamus volume (left panel). This effect was dose-dependent (central panel) and partially replicated in an independent adolescent cohort (right panel).

**Table 2.**
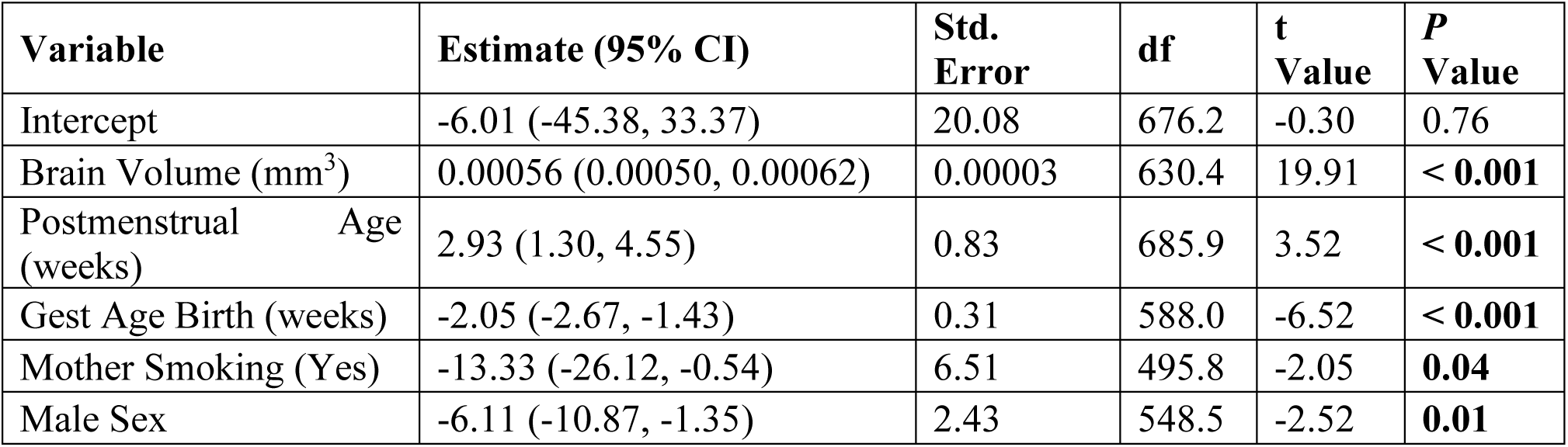
Linear Mixed-Effects Model Predicting Whole Hypothalamus Volume *(N = 699 observations from 631 participants)*.

The associations between infant hypothalamus volume and GA, sex, and maternal smoking survived (and often improved in effect size and significance) exhaustive testing in the context of different models (ratio vs. non-ratio), global adjustment covariates (brain volume, bilateral amygdala volume), and subsets of data with more narrow limits (PMA 40-43 weeks, GA 39-41 weeks, near full-term >36 weeks). Such exhaustive testing was used to minimize the confounding between GA and PMA at scan present in the dHCP dataset (see *Supplementary Materials S1: Multiverse Analyses*).

### 3.4 Maternal Smoking During Pregnancy Is Associated with Adolescent Hypothalamus Volume

Effects of adolescent age, sex, preterm birth status, and maternal smoking during pregnancy on hypothalamus volume were considered in the ABCD sample. Adolescent age was associated with hypothalamus volume in both absolute and adjusted measures (t_age,abs_=37.6, p_age,abs_<10^-10^; t_age,adj_=39.9, p_age,adj_<10^-10^) in a direction supportive of protracted growth during the adolescent period, including over and above global volume. Absolute and relative measures of hypothalamus volume were associated with sex (males 1.1% larger on average; t_sex,abs_=-10.2, p_sex,abs_<10^-10^; t_sex,adj_=-2.3, p_sex,adj_=0.02), and sex-specific associations with age were significant in both absolute and relative terms (t_age-sex,abs_=-5.6, p_age-sex,abs_<10^-7^; t_age-sex,rel_=-5.3, p_age-sex,rel_<10^-6^). Collectively, these findings suggest that the hypothalamus during early adolescence is characterized by sex-specific growth trajectories in which the male hypothalamus volume is larger than in females and increasing in magnitude. There was no observed main (t=-0.9, p=0.39) or sex-specific (t=0.7, p=0.47) association with preterm birth status. However, there was a significant main effect of maternal smoking during pregnancy (t_smoking_=-3.0, p_smoking_^=^0.002) that survived adjustment for potentially confounding variables (t_smoking_=-2.8, p_smoking_^=^0.005). Consistent with observations in the dHCP dataset, our result suggested a similar reduction in adolescent hypothalamus volume (-1.5% on average) in the presence of maternal smoking during pregnancy (**Figure 4**). There was no significant interaction effect between maternal smoking during pregnancy and offspring sex (t=0.6, p=0.54).

## 4. Discussion and Conclusions

Using large, well-characterized neuroimaging datasets, we demonstrate that prenatal factors, such as GA, PMA at MRI, sex, and maternal smoking during pregnancy, are associated with offspring hypothalamus volume. These findings highlight four key observations. First, although raw, unadjusted hypothalamus volume was positively associated with PMA, this association was no longer present when controlling for global BV, either through covariate adjustment or ratio scaling. Second, GA was negatively associated with adjusted hypothalamus volume; however, evidence from adolescence suggests that any lasting impact of preterm birth on hypothalamus volume is subtle or undetectable. Third, our findings suggested sexual dimorphism in absolute and relative hypothalamus volume and evidence of sex-specific effects on the associations between volume and age measures across development. Finally, maternal smoking during pregnancy was linked to smaller hypothalamus volume at birth, and this association was partially replicated in a large-scale, nationally representative adolescent sample in both magnitude and direction of effect.

Unadjusted analyses from the dHCP newborn dataset revealed a linear increase in hypothalamus and brain volumes with PMA, with BV having a steeper slope. However, this association was no longer present when adjusting for BV, providing mixed evidence that hypothalamic growth during this early life period may reflect global (or local) expansion. In adolescence, however, hypothalamus volume was positively associated with age in both absolute and BV-adjusted terms. Notably, these observations are consistent with prior literature (37) in smaller samples (n=26, 0-6 mos; n=43, 10-12 yrs) demonstrating no significant increase in hypothalamus volume relative to BV in the early life period, yet a significant increase in hypothalamus volume relative to BV in the adolescent period. Together, these data support a model in which hypothalamic growth broadly parallels global brain development, yet exhibits distinct, phase-specific acceleration patterns likely reflecting the structure’s responsiveness to age-related physiological changes such as puberty.

We observed that lower GA at birth was associated with larger hypothalamus volume after adjusting for PMA and BV. This relationship may reflect early adaptation or accelerated hypothalamic development in response to the extrauterine environment. One possible explanation is that preterm birth acts as a developmental catalyst, triggering early physiological transitions, including hormonal, metabolic, and neurodevelopmental changes that transiently accelerate hypothalamic growth, analogous to increased functional connectivity in stress-related circuits in premature infants (38). Interestingly, prior work suggests that energy homeostasis-relevant hypothalamic *subunit* growth in preterm infants may diverge early, potentially contributing to smaller volumes observed in adults born preterm (23). However, it should be noted that the evidence presented here suggests that the impact of preterm birth on whole hypothalamus volume is no longer detectable during adolescence. Taken together with the findings from Ruzok et al., we speculate that the impact of preterm birth on hypothalamus structure persists across the lifespan but is masked by age-related physiological changes such as puberty, and/or is subunit specific.

We identified multiple sex-specific effects in hypothalamic development, including both volumetric differences and distinct associations between hypothalamus volume and age. In newborns, we observed that the hypothalamus is larger in males than in females on average. Our finding is consistent with an abundance of literature supporting this premise (39–42). However, when adjusting for global brain measures, the female hypothalamus was observed to be larger. Notably, this finding is consistent with recent literature demonstrating a similar effect in the newborn caudate nucleus using a large-scale multi-site secondary analysis (43), and newborn total gray matter volumes from the same cohort considered here (but with no measures for the hypothalamus available (40). Such relative differences may stem from differential timing of regional brain growth as a function of age and sex, likely driven by complex physiological changes and needs (44–47). In contrast, by adolescence, we found that the hypothalamus was larger in males than in females, both in absolute and relative terms. These findings motivate future research examining how developmental timing, hormonal regulation, and energetic needs influence hypothalamic growth trajectories.

In addition to sex-based differences in hypothalamus volume, we also observed sex-specific associations between volume and age. Specifically, relative to global brain development across GA, the hypothalamus grows at a slower rate, with males affected to a greater extent than females. Due to the absence of any functional or behavioral measurements, the implications of such a slower relative growth are unclear at this point. Our study provides an opportunity to further validate the current findings and add behavioral or functional effects to understand the clinical relevance of the relatively slower hypothalamic growth rate in males than females.

The reduced hypothalamus volume in newborns and adolescents whose mothers reported smoking during pregnancy, along with the dose-dependence of this relationship in newborns, suggests a direct adverse impact of maternal smoking on hypothalamic development. Nicotine and other neuroactive compounds in cigarette smoke readily cross the placenta, exposing the rapidly developing fetal brain to harmful toxins (48). Nicotine acts by binding to nicotinic acetylcholine receptors (nAChRs), which are critical for typical neurodevelopmental processes, including neurogenesis, neuronal migration, and synaptogenesis (49,50). In utero exposure to nicotine can interfere with neurotransmitter signaling and disrupt these fundamental cellular processes (49,51). It is unclear why hypothalamus, but not amygdala, volume was reduced with nicotine exposure. While both structures express nAChRs and are, therefore, expected to be similarly susceptible to the nicotine effects, the differential susceptibility may be attributed to the differences in developmental trajectories (52,53), critical periods for cellular events, the precise subtypes of nAChRs expressed in each region (54,55), and the timing of prenatal exposure relative to these sensitive windows.

Limitations of our study include confounding between age measures (PMA and GA) in early life, simplification of the hypothalamus as a whole structure, and largely cross-sectional measures. The dHCP study collected newborn MRI at less than 3 postnatal weeks on average (median less than two weeks), meaning that PMA is partially confounded with GA. To address this confounding effect, we have considered variance inflation factors where necessary and provided sensitivity analyses using subgroups to minimize confounding and reaffirm hypotheses.

However, findings centered on the effects of GA should still be interpreted with caution, and future replication studies are warranted. The hypothalamus consists of a heterogeneous and specialized nuclei structure. Therefore, such heterogeneity is often oversimplified when considering the hypothalamus as a whole. Thus, while outside the scope of our current investigation, future efforts will aim to parcellate the newborn hypothalamus into more structurally and functionally informed subunits (56). Finally, this study has attempted to consider the structure of the developing hypothalamus using two large-scale publicly available datasets. However, there exists a large age gap and an obvious lack of longitudinal data between studies. Given a strong body of preclinical literature supporting prenatal effects on the developing hypothalamus, the clinical evidence provided here, and the promise of the forthcoming Healthy Brain and Child Development (HBCD) study (57), we assert that there exists a unique opportunity to test and extend our hypotheses by incorporating longitudinal trajectories, biological mechanisms, and behavioral outcomes to further understand the multisystemic effects of this small, but important brain region.

The current study highlights the feasibility of studying the MRI-based structure of the developing hypothalamus and presents new insights into how perinatal factors may shape life course outcomes through their effects on this “small and mighty” (11) brain region. Our study advances the premise that hypothalamic development is shaped by gestational age at birth, maternal smoking during pregnancy, and provides further evidence for sex-specific programming and developmental effects. Using two large developmental cohorts, our study provided robust evidence that the hypothalamus is a developmentally sensitive and underexplored target of perinatal influences. Given the hypothalamus’ central role in coordinating metabolic, endocrine, and behavioral systems vital for early survival and lifelong adaptation, understanding its development is essential for uncovering how early life exposures shape trajectories of health and disease.

## Data and Code Availability Statement

All underlying data are openly available upon request. Hypothalamic segmentations and code will be made available privately upon request during the review process and publicly available following publication (https://github.com/jerodras).

## Acknowledgements

We thank the participant volunteers for donating their time to the dHCP and ABCD projects and the generosity of the European Research Council and the National Institute of Health for making the dHCP and ABCD data publicly available.

## Prior Presentation

This work has not been presented prior to submission.

