## Supplement File for "MRI-Based Structural Development of the Human Newborn Hypothalamus"

***Supplementary Materials S1: Multiverse Analysis***

Models from the main manuscript were repeated using various design choices in a multiverse analysis. Two considerations were made: 1) the variable used for global volume adjustment, and 2) the sample used with constraints on participant filtering.

For global adjustment, four separate models were run. First, we repeated the model used in the main manuscript as a reference model. This model included total brain volume as a covariate in the model to account for variance in total size, thereby specifying effects to hypothalamus volume. Second, the reference model was repeated substituting total amygdala volume as a subcortical structure similar in size and proximity to the hypothalamus. Third, the reference model was repeated substituting total intracranial volume (ICV, another commonly used proxy for brain size) for brain volume. Last, the outcome was changed to a ratio of hypothalamus to brain volume. Importantly, this ratio measure helps alleviate concerns about non-linearities not captured in linearly adjusting for total size.

For sample adjustment, we identified four subsamples to help deepen understanding of the observed effects. First, we repeated the models using an imaged-post-term (postmenstrual age at scan, PMA, >40 weeks) sample to address concerns about partial voluming in especially young newborns. Second, we repeated the models using a narrow gestational age window (39-41 weeks) to minimize variance and effects of gestational age at birth. We note that this subsample is *a priori* expected to have small gestational age (GA) of birth effect sizes. Third, we repeated the models using a full term-born sample to minimize effects of pre-term birth. While we believe this lack of variance is not ideal for primary hypothesis testing, it may help inform on effects independent of unobserved clinical and biological confounds from pre-term birth. Last, we repeated the models using a sample with a narrow (2-4 days) postnatal scan age (SA) to help isolate effects of gestational age at birth from postmenstrual age at scan. Models using this last sample excluded the postmenstrual age at scan term due to its high correlation with gestational age at birth, by design.

Finally, each of these eight total models were repeated for each of the primary findings in the main manuscript: the main effect of gestational age at birth, the interaction between sex and gestational age at birth, and maternal smoking during pregnancy. Results of these models are discussed in line with their respective multiverse tables.

***Supplementary Table S1: Main Effect of Gestational Age at Birth***

Model Specification in R: *lmer(hypoth_vol ~* ***size +*** *pma +* ***ga*** *+ sex + ( 1 | participant_id)^*^*

| **Sample** | **n(Observ.)** | **Adjustment** | **Beta-std** | **t-value** | **p-value** |
| --- | --- | --- | --- | --- | --- |
| *Full (Ref.)* | *699* | *BV (Ref. Model)* | *-0.144 [-0.19 -0.10]* | *-6.42* | *<10⁻¹⁰* |
| *Full* | *699* | *Amyg* | *-0.217* | *-8.39* | *<10⁻¹⁰* |
| *Full* | *699* | *ICV* | *-0.143* | *-6.37* | *<10⁻¹⁰* |
| *Full* | *699* | *Ratio* | *-0.280* | *-5.88* | *<10⁻¹⁰* |
| *PMA>40wks* | *288* | *BV* | *-0.239* | *-5.84* | *<10⁻^7^* |
| GA=39-41wks | 227 | BV | 0.011 | 0.24 | 0.814 |
| GA >36wks | 468 | BV | -0.081 | -1.88 | 0.061 |
| *SA=2-4days* | *168* | *BV* | *-0.204* | *-2.74* | *0.007* |

*^*^Linear mixed effects models were used when repeated measures were present, whereas simple linear regression (lm) models were used with sub-samples composed only of single measures.*

Five of the seven alternate models supported a robust (survived significance) effect of gestational age at birth. The full-term birth sample, while consistent in direction and magnitude given the reduced variance in GA, failed to achieve significance at a p<0.05 threshold. The last was constrained to a narrow GA (39-41 weeks) window and was *a priori* not expected to demonstrate an effect of GA. Notably, three of the alternate models had effect sizes outside (greater than) the confidence interval observed in the reference model. Collectively, these analyses support an effect of gestational age at birth that is robust to model choice and design.

***Supplementary Table S2: Sex-Specific Effects of Gestational Age at Birth***

Model Specification in R: *lmer(hypoth_vol ~* ***size +*** *pma +* ***ga x sex*** *+ ( 1 | participant_id)^*^*

| **Sample** | **n(Observ.)** | **Adjustment** | **Beta-std** | **t-value** | **p-value** |
| --- | --- | --- | --- | --- | --- |
| *Full (Ref.)* | *699* | *BV (Ref. Model)* | *-0.329 [-0.60 -0.06]* | *-2.36* | *0.019* |
| Full | 699 | Amyg | -0.284 | -1.70 | 0.090 |
| *Full* | *699* | *ICV* | *-0.322* | *-2.30* | *0.022* |
| *Full* | *699* | *Ratio* | *-0.598* | *-2.01* | *0.045* |
| *PMA=40-43wks* | *288* | *BV* | *-1.098* | *-2.86* | *0.005* |
| GA=39-41wks | 227 | BV | -4.721 | -1.47 | 0.142 |
| *GA >36wks* | *468* | *BV* | *-1.835* | *-2.27* | *0.024* |
| SA=2-4days | 168 | BV | -0.254 | -0.26 | 0.796 |

*^*^Linear mixed effects models were used when repeated measures were present, whereas simple linear regression (lm) models were used with sub-samples composed only of single measures.*

Four of the seven alternate models supported a robust (survived significance) interaction effect of gestational age at birth by sex. The model adjusting for amygdala volume, while consistent in direction and magnitude (within confidence interval), failed to achieve significance at a p<0.05 threshold. The model isolating GA by limiting SA (2-4 days) failed to replicate the interaction effect. The last model not replicating the finding was constrained to a narrow GA (39-41 weeks) window and was *a priori* not expected to demonstrate an effect of GA. Collectively, these analyses support sex-specific effects of gestational age at birth but also suggest model selection may be critical for their observation.

***Supplementary Table S3: Main Effect of Maternal Smoking Status During Pregnancy***

Model Specification in R: *lmer(hypoth_vol ~* ***size +*** *pma + ga + sex +* ***smoking*** *+ ( 1 | participant_id)^*^*

| **Sample** | **n(Observ.)** | **n(Smoking)** | **Adjustment** | **Beta-std** | **t-value** | **p-value** |
| --- | --- | --- | --- | --- | --- | --- |
| *Full (Ref.)* | *699* | *23* | *BV (Ref. Model)* | *-0.036 [-0.07 -0.00]* | *-2.05* | *0.041* |
| *Full* | *699* | *23* | *Amyg* | *-0.053* | *-2.59* | *0.009* |
| *Full* | *699* | *23* | *ICV* | *-0.037* | *-2.12* | *0.035* |
| *Full* | *699* | *23* | *Ratio* | *-0.083* | *-2.23* | *0.026* |
| PMA=40-43wks | 288 | 8 | BV | -0.033 | -0.82 | 0.412 |
| GA=39-41wks | 227 | 7 | BV | -0.060 | -1.43 | 0.155 |
| GA >36wks | 468 | 10 | BV | -0.045 | -1.49 | 0.137 |
| SA=2-4days | 168 | 6 | BV | -0.096 | -1.86 | 0.064 |

*^*^Linear mixed effects models were used when repeated measures were present, whereas simple linear regression (lm) models were used with sub-samples composed only of single measures.*

Three of the seven alternate models supported a robust (survived significance) effect of maternal smoking status at birth. However, the four that did not were statistically underpowered due to subsampling. Of the three that did survive significance, all were of larger effect size though only one (ratio model) was outside the confidence interval of the reference model. Of the four underpowered models, while difficult to interpret with confidence, all were consistent in direction and magnitude of effect. Collectively, these analyses support an effect of maternal smoking status that is robust to model choice and design.
